# Osmotic stress via calmodulin lead to the formation of stress granule in Drosophila S2 cells

**DOI:** 10.1101/2022.03.28.485805

**Authors:** Chujun Zhang, Rianne Grond, J. Mirjam A. Damen, Wei Wu, Catherine Rabouille

## Abstract

Cellular stress of S2 cells leads to the formation of stress assemblies by phase separation of cytoplasmic components. We have shown that the cellular stresses of either high increase of the NaCl concentration in the extracellular medium, or a moderate one combined to amino acid starvation, leads to the formation of Sec bodies where components of the endoplasmic Reticulum exit sites (ERES) coalesce. These extracellular stresses lead to both the activation of salt inducible kinase (SIK), and to ER stress triggering the activation of the two downstream kinases IRE1 and PERK. Interestingly, the same stresses also result in the formation of a second stress assembly, the stress granules, which stores specific RNAs and RNA binding proteins. Here we asked whether stress granule formation is governed by the same pathways as Sec bodies. However, we found that the inhibition of SIK, IRE1 and PERK does not affect stress granule formation. Instead, we found that osmotic stress through the addition of either salts (including calcium chloride) or sucrose leads to the formation of stress granules. Interestingly, stress granule formation is partly modulated by calmodulin activation, suggesting the involvement of calcium signaling. Furthermore, as Sec body formation is driven by entirely different pathways, these results show that the same cells under the same stress, form two different stress assemblies by non-overlapping downstream pathway activation, perhaps explaining that they do not coalescence into a single structure.

## Introduction

There are two forms of cell compartmentalization: membrane-bound organelles and membrane-less organelles (MLOs). Unlike membrane-bound organelles, MLOs are not surrounded by a sealed phospholipid membrane and are formed by phase separation (Banani et al., 2016; Gomes and Shorter, 2018; Patel et al., 2015). Phase separation drives a homogeneous solution of dispersed macromolecules into two distinct phase which can be either liquid or solid. MLOs (such as nucleolus) exist in cells in basal conditions, but it is now well established that cellular stress leads to phase separation and the formation of many different MLOs (van Leeuwen and Rabouille, 2019), including stress granules (Anderson and Kedersha, 2002; Aulas et al., 2017) and Sec bodies (Zacharogianni et al., 2014).

Sec bodies are stress assemblies that first have been identified in Drosophila S2 cells under the condition of amino acid starvation in the Krebs Ringer Bicarbonate buffer, KRB. KRB incubation results in the remodeling and coalescence of ER exit site (ERES) components into round, membrane-less, reversible, liquid-like, Sec bodies of 0.6-0.8 μm in diameter (Zacharogianni et al., 2014). KRB contains a 2.6-fold higher NaCl concentration than the growing medium Schneider’s and does not contain amino acids. Interestingly, a 4-fold increase in NaCl concentration in Schneider’s (through the specific addition of 150mM NaCl, SCH150) also leads to Sec body formation in S2 cells (Zhang et al., 2021).

This led us to identify signaling pathways necessary for Sec body formation. The first one is the activation of Salt Inducible Kinases (SIKs) by NaCl, which is sufficient when the NaCl stress is high, for instance in SCH150. The second is the activation of IRE1 and PERK downstream of ER stress (Walter and Ron, 2011) that is induced by amino acid starvation (Zhang et al., 2021). IRE1 and PERK activation in solo does not lead to Sec body formation. It needs to be combined with moderate salt stress, as in KRB. Accordingly, addition to dithiothreitol (DTT that activates both IRE1 PERK) together with 100mM NaCl to Schneider’s leads to the same efficient Sec body formation as KRB incubation (Zhang et al., 2021).

We have reported that upon KRB incubation, not only Sec body formed, but also stress granules in close proximity and with a similar kinetics (Aguilera-Gomez and Rabouille, 2017; Zacharogianni et al., 2014). Stress granule is another class membrane-less and reversible MLOs that has been widely reported to contain mRNAs bound to RNA-binding proteins (Anderson and Kedersha, 2002). The RNAs that phase separated into stress granules are stored there during the stress and can be immediately translated upon stress relief. In Drosophila S2 cells, they can be visualized by labeling several RNA binding proteins, such as FMR1 (Zacharogianni et al., 2014), Caprin and Rasputin (Aguilera-Gomez et al., 2017). Most of the stress granules formation is dependent on eIF2alpha phosphorylation resulting in translational stalling (canonical pathways). However, they can also form in an independent manner (non-canonical pathways) like those activated upon osmotic stress (Aulas et al., 2017; Kedersha et al., 2016). In this regard, in our study, we show that stress granule formation is also induced by incubating cells in SCH150, i.e., addition of NaCl to the growing medium (**Figure 1A**).

**Figure 1:**
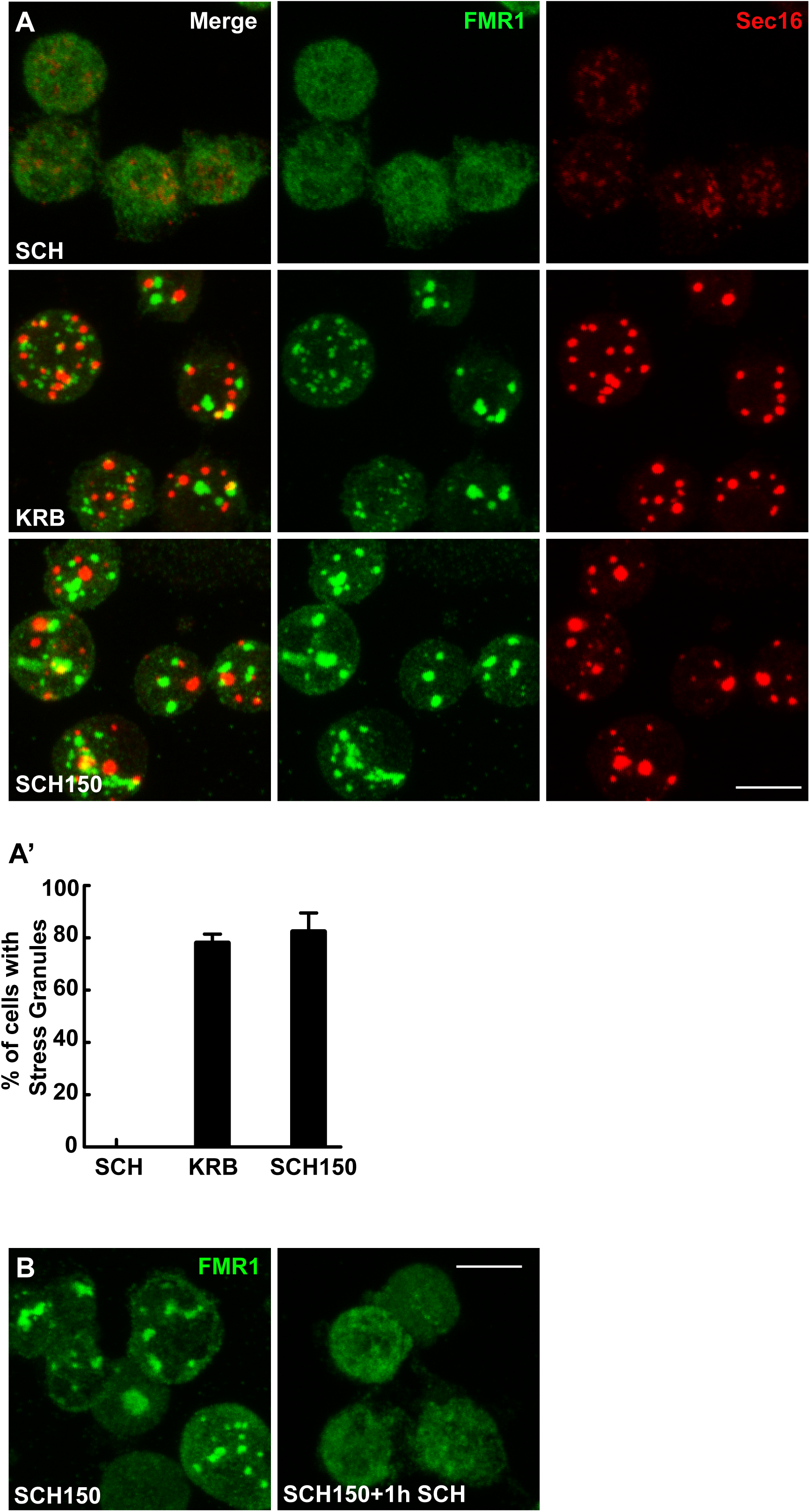
KRB and SCH150 incubation leads to reversible stress granule formation. (A, A’) Immunofluorescence (IF) visualization of endogenous FMR1 and Sec16 in S2 cells in growing medium Schneider’s (SCH) and in cells incubated in KRB and SCH150 (addition of 150mM of NaCl to Schneider’s) for 4 h at 26°C (A). Note that, upon KRB and SCH150 incubation FMR1 and Sec16 remodel into two distinct large structures, the stress granules and Sec bodies, respectively ((A). Quantification of stress granule formation in KRB and SCH150 (A’). (B) IF visualization of stress granule formation (marked by FMR1) in SCH150 for 4 h at 26°C, as well as SCH150 for 4 h followed by 1h SCH incubation at 26°C, showing their reversibility. Scale bars: 10µm. Errors bars: SEM.

Here, we asked whether, stress granule formation in response to KRB and SCH150 is governed by the activation of the same kinases as Sec bodies. However, we found that the inhibition of SIK, IRE1 and PERK in solo does not inhibit stress granule formation upon KRB and SCH150. Instead, we found that the addition of several different salts (including CaCl2) to the growing medium Schneider’s as well as osmotic stress (by addition of sucrose) is enough to lead to stress granule formation. This is strikingly different than for Sec bodies, which formation strictly depends in the specific presence and the addition of NaCl and do not form upon addition of sucrose (Zhang et al., 2021). Mass spectrometry reveals that calmodulin is one of the most upregulated proteins during KRB incubation, and we found that its depletion partially prevents stress granule formation both in KRB and SCH150. This suggests that calcium signaling plays a role in stress granule formation, in line with the presence of calcium binding proteins in these structures (Markmiller et al., 2019; Marmor-Kollet et al., 2020). Furthermore, these results show that the same cells, under the same stresses form two different stress assemblies by non-overlapping downstream kinase activation.

## Results

### KRB and high NaCl stress leads to stress granule formation

As mentioned in the introduction, the same cellular stresses (KRB and SCH150 for 4h) of S2 cells not only lead to the formation of Sec body (Zacharogianni et al., 2014; Zhang et al., 2021), but also of stress granule that are marked here by FMR1 (**Figure 1A, A’**). We have shown that the stress granules formed upon KRB appears to be *bona fide* stress granules, as in addition to FMR1, they contain several RNA binding proteins and are membraneless and are reversible (Aguilera-Gomez et al., 2017). Here, we confirmed that SCH150 induced stress granules are reversible. Indeed, they dissolved upon furthermore incubation in growing medium for 1h following the stress period (**Figure 1B**). Furthermore, they contain Rox8, the homolog of TIA1 in mammalian cells (Ma et al., 2017), a key stress granule marker in mammalian cells (**Figure 4D**). Furthermore, stress granules form around poly-adenylated (polyA) mRNAs that are stored in these structures during the period of stress(Kedersha et al., 1999). Here, we showed the stress granule formed in KRB and SCH150 contain polyA mRNAs using fluorescence in situ hybridization (FISH) with an oligo (dT) probe (**Figure 5**). Taken together, these results indicate that the stress granules formed in high salt stress (SCH150) are *bona fide* stress granules.

Given that the same stress leads to the simultaneous formation of two distinct stress assemblies (**Figure 1**), it leads us to question whether stress granule formation follow the same pathways and are governed by the same activation of the same kinases as Sec bodies.

### SIK and three ER stress sensors are not involved in stress granule formation upon KRB and SCH150

As mentioned above, we have shown that SIK activation is an important contributor in Sec body formation. To answer whether SIK is activation also plays a role in stress granule formation, we first monitored their formation in cells incubated in SIK inhibitor HG-9-91-01, both upon KRB and SCH50. However, this inhibitor has no effect on stress granule formation (**Figure 2A**), suggesting that SIK is not involved in stress granule formation, neither in KRB and SCH150.

**Figure 2:**
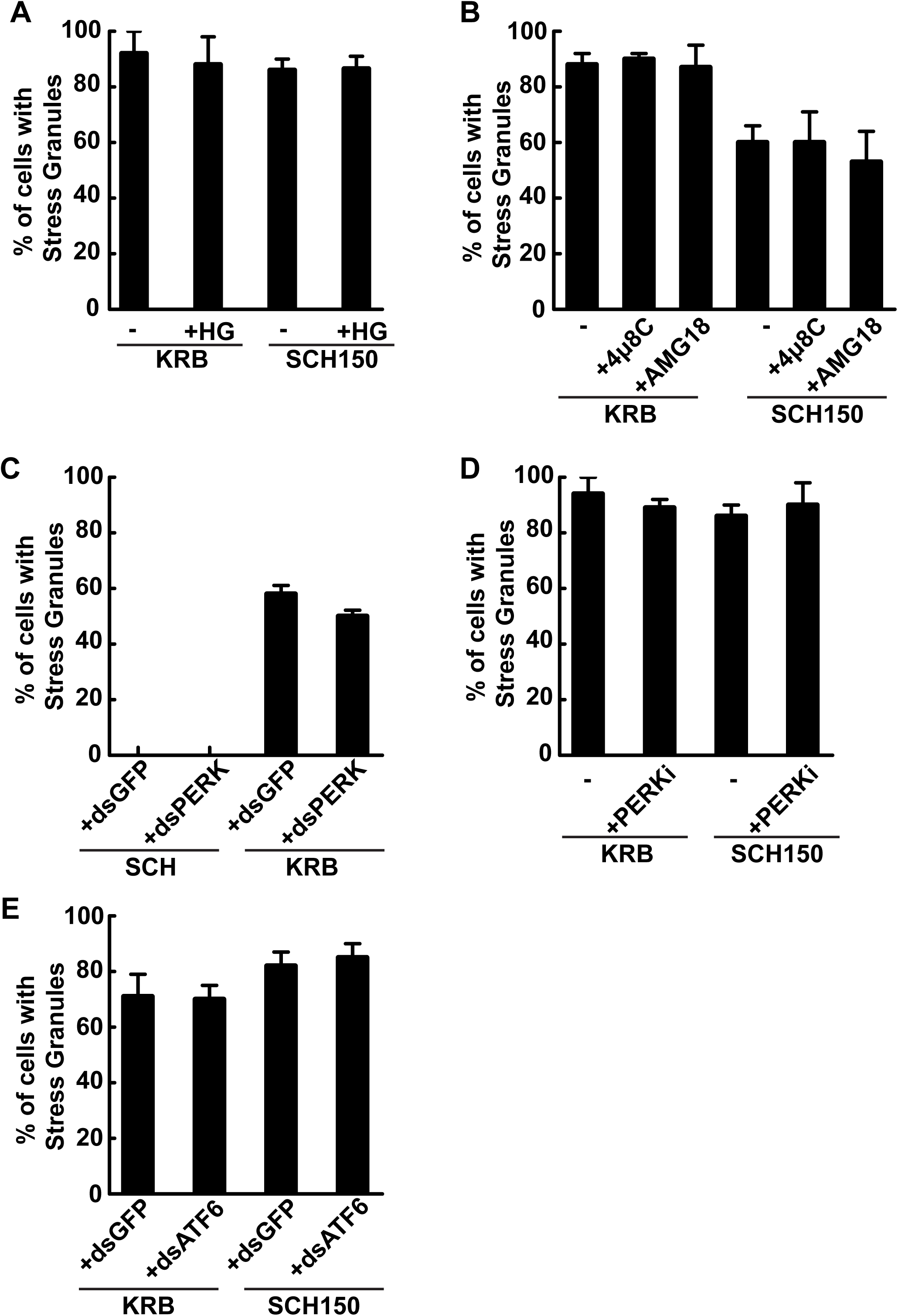
SIK and ER stress kinases are not involved in KRB and SCH150 induced stress granule formation. (A) Quantification of stress granule formation (marked by FMR1) in cells incubated in KRB and SCH150 with or without the SIK inhibitor HG-9-91-01 (5µM). (B) Quantification of stress granule formation (marked by FMR1) in cells incubated in KRB and SCH150 with or without the IRE1 RNAse inhibitor 4µ8C (30µM) and IRE1 kinase inhibitor AMG-18 (10µM). (C) Quantification of stress granule formation (marked by FMR1) in cells upon mock depletion (dsGFP) or PERK depletion (dsPERK) incubated in KRB. (D) Quantification of stress granule formation (marked by FMR1) in cells incubated in KRB and SCH150 with or without the PERK inhibitor (5µM). (E) Quantification of stress granule formation (marked by FMR1) in cells upon mock depletion (dsGFP) or ATF6 depletion (dsATF6) incubated in KRB and SCH150. Errors bars: SEM

KRB and SCH150 incubation leads to an increase of the protein level of the ER chaperone Bip that we take as a measure of the UPR stimulation by ER stress (Zhang et al., 2021). UPR activation is downstream of the 3 kinases that are ER stress sensors IRE1, PERK and ATF6. Those can be activated upon Bip release from their luminal domain during stress (Hetz, 2012). In the case of IRE1, both its kinase and RNAse domains are activated. In order to investigate the role of IRE1 activation in stress granule formation in KRB and SCH150, we incubated the cells with AMG-18 (IRE1 kinase inhibitor) and 4µ8C (IRE1 RNAse inhibitor) under conditions of stress (KRB and SCH150). However, neither inhibitor reduce stress granule formation (**Figure 2B**), whereas Sec body formation was strongly inhibited showing that both drugs are efficient in Drosophila cells (Zhang et al., 2021). This suggests that IRE1 activation does not play a role in stress granule formation in this system. We then tested the role of PERK and ATF6 using depletion by RNAi. Again, KRB or SCH150 induced stress granule formation was not reduced (**Figure 2C, E**). Furthermore, the PERK inhibitor GSK2606414 did not have any effect on stress granule formation upon KRB and SCH150 (**Figure 2D**). This suggested that PERK and ATF6 activation are not necessary for stress granule formation.

Taken all these together, we conclude that the activation of neither SIK, nor any of the three ER stress sensors (IRE1, PERK and ATF6) are involved in stress granule formation that appears to be mediated by the activation of different kinases than Sec bodies.

### The amino-acid starvation sensor GCN2 is not involved in the KRB and SCH150 induced stress granule formation

We had shown previously that RNA translation is inhibited in cells incubated in KRB (Aguilera-Gomez et al., 2017). This is characterized by the phosphorylation of eIf2alpha (Hetz, 2012). In this regard, we show that eIf2alpha is phosphorylated in S2 cells incubated in both KRB and SCH150 (**Figure 3A**). In Drosophila, eIf2alpha phosphorylation is typically mediated by PERK and GCN2 (General Control Nonderepressible 2). Interestingly, GCN2 is also known as an amino acid starvation sensor (Dever and Hinnebusch, 2005; Towle, 2007) and this might be relevant to the KRB incubation as it does not contain amino-acids. To test whether GCN2 plays a role in KRB and SCH150 induced stress granule formation, we inhibited GCN2 with a specific inhibitor (GCN2iB). However, stress granule still forms as in control conditions (absence of inhibitors) (**Figure 3B**) suggesting that the activation of GCN2 is not involved in stress granule formation either in KRB or SCH150.

**Figure 3:**
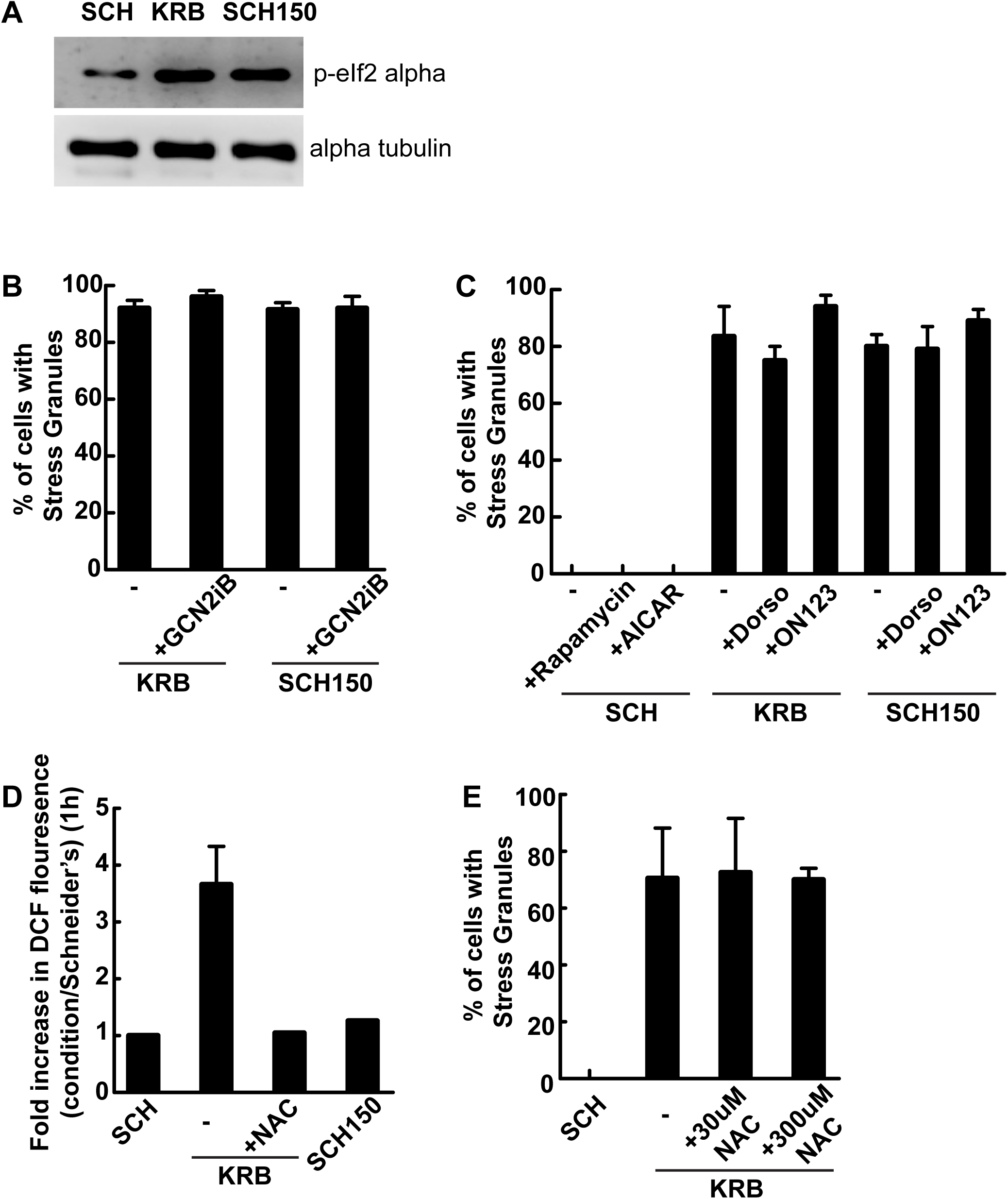
GCN2, mTORC1, AMPK and ROS production are not involved KRB and SCH150 induced stress granule formation. (A) Western blot visualization of eIF2alpha phosphorylation in cells in SCH, KRB and SCH150. (B) Quantification of stress granule formation (marked by FMR1) in cells incubated in KRB and SCH150 with or without GCN2 inhibitor GCN2iB (10µM). (C) Quantification of stress granule formation (marked by FMR1) in cells incubated in SCH with or without AICAR (1mM) and Rapamycin (10µM), and cells incubated in KRB and SCH150 with or without Dorsomorphin (Dorso, 1µM) and ON123300 (ON123, 10µM). (D) Quantification of the increase in DCF fluorescence measuring the production of ROS upon KRB, KRB with addition of ROS inhibitor N-acetyl-L-cysteine (NAC) and SCH150 when compared to SCH. (E) Quantification of stress granule formation (marked by FMR1) in cells incubated in SCH, KRB, KRB with addition of N-acetyl-L-cysteine (NAC, 30µM or 300µM). Errors bars: SEM.

### Nutrient sensors pathways mTORC1 and AMPK activation are not involved in stress granule formation upon KRB and SCH150

mTORC1 (mechanistic target of rapamycin complex I) activation is also a sensor of the presence of circulating amino acids in the medium (Kim and Guan, 2019; Manifava et al., 2016). It is inhibited when amino-acid level is low in the circulating medium(Condon and Sabatini, 2019). Ribosomal S6 kinase is a direct target of mTORC1, mTORC1 phosphorylates S6 upon its activation (Jacinto et al., 2004; Li et al., 2010). We showed that S6 no longer phosphorylated after KRB incubation (Zhang et al., 2021). Interestingly, the absence of S6 phosphorylation also seal RNA translation, which could lead to stress granule formation. In this regard, we tested whether the sole inhibition of mTORC1 (using Rapamycin) would lead to stress granule formation (Zhang et al., 2021). However, the sole inhibition of mTORC1 by rapamycin did not trigger stress granule formation (**Figure 3C**). This suggested that amino-acid starvation inhibits mTORC1, but only inhibiting mTORC1 is not enough to trigger stress granule formation. Taken together, these results indicate that the KRB and SCH150 induced stress granule formation in S2 cells is not linked to translation inhibition, as neither GCN2 nor PERK activation (**Figure 2**) nor does mTORC1 inhibition modulate their formation.

AMPK (AMP-activated protein kinase) is a cellular energy sensor and activated by a decrease in intracellular ATP (Miyamoto et al., 2008; Perera and Turner, 2015). As we observed that a sharp decrease in the intracellular concentration of ATP correlates with Sec body formation in KRB (Zhang et al., 2021), we examined whether AMPK activation is involved in stress granule formation in KRB. To test this, we first incubated S2 cells with AMPK activator AICAR (Ducommun et al., 2014) in growing conditions but this did not lead to stress granule formation (**Figure 3C**). Furthermore, we tested whether an inhibitor of AMPK had an inhibitory effect on KRB and SCH150 induced stress granule formation. However, neither Dorsomorphin (Weiss et al., 2010), nor ON123300 (Zhang et al., 2021) showed an effect of stress granule formation (**Figure 3C**). These results suggest that AMPK activation is not necessary for stress granule formation.

Taken together, the above results indicate that mTOR1 and AMPK are not involved in stress granule formation.

### ROS production is not necessary for stress granule formation

To assess which other pathways are stimulated upon KRB incubation that could lead to stress granule formation, we had previously performed an RNA sequencing experiment, the analysis of which suggested that KRB elicits oxidative stress (Zhang et al., 2021). To complete this approach, we performed an analysis by mass spectrometry to identify proteins that are specifically enriched or depleted in cells incubated in this buffer. Overall, KRB incubation for 4h leads to a 30% reduction in protein level, but 59 proteins appear enriched in KRB and 94 appear depleted (Suppl Table S1). In particular, the level of the oxidative stress markers peroxiredoxins (Poynton and Hampton, 2014) were reduced after 2h of KRB incubation (not shown), a result in agreement with the RNA sequencing (Zhang et al., 2021). Taken together, the KRB protein and gene signature is suggestive of oxidative stress, indicating that KRB incubation elicits ROS production.

We have previously used DCF fluorescence measurement to visualize ROS production upon KRB and showed that it is efficiently inhibited by N-acetyl-L-cysteine (NAC) (Zhang et al., 2021). In this regard, we tested whether ROS inhibition prevents stress granule formation. However, ROS production inhibition did not modify KRB and SCH150 induced stress granule formation (**Figure 3D**). This is in line with the fact that SCH150 did not elicit ROS (**Figure 3E**). Taken together, ROS production is not necessary for stress granule formation.

### Osmotic stress leads to stress granule formation

We then focused on the non-conventional pathways leading to stress granule formation (see introduction) (Aulas et al., 2017; van Leeuwen and Rabouille, 2019), including osmotic stress. As SCH150 corresponds to a 4-fold increase of Na+ in the cell incubation medium (2.6-fold for KRB), we tested whether these media led to an increase of cytoplasmic Na+ concentration that can be visualized using Sodium Green Indicator a visible light excitable probe. As expected, we found an Na+ concentration increase in KRB and SCH150 (**Figure 4A**).

**Figure 4:**
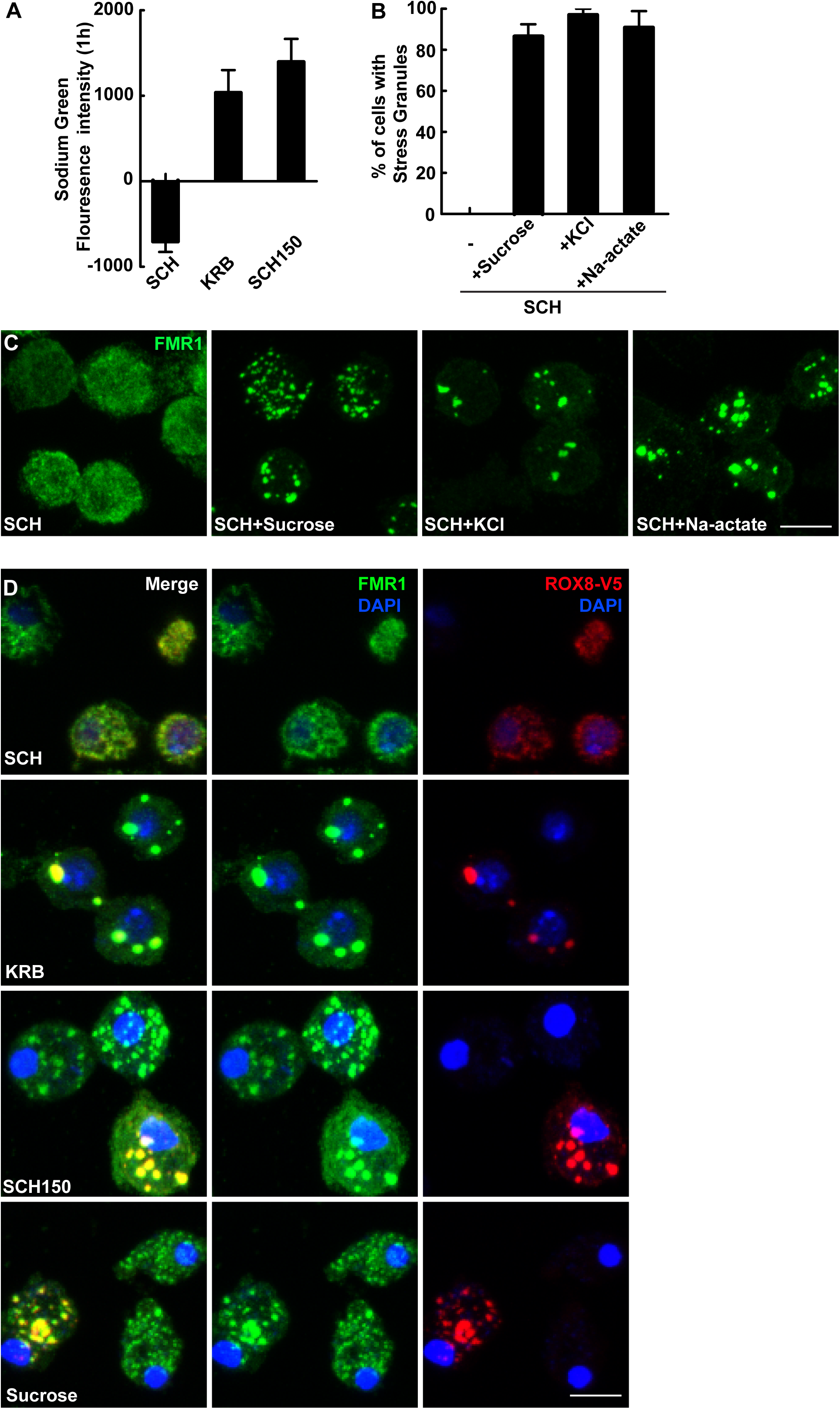
Osmotic stress leads to stress granule formation. (A) Quantification of the increase in Sodium Green fluorescence intensity (measuring the intracellular Na^+^ level) upon SCH, KRB and SCH150. (B, C) Quantification (B) and IF visualization (C) of stress granule formation (marked by FMR1) in SCH supplemented with sucrose (400mM), KCl (150mM) or Na-acetate (150mM) for 4h at 26°C. (D) IF visualization of overexpressed Rox8-V5 (V5, red) and FMR1 (green) in cells incubated in SCH, KRB, SCH150 and SCH+sucrose (400mM). Note that FMR1 positive stress granules also contain Rox8-V5. Scale bars: 10µm. Errors bars: SEM.

Accordingly, we asked whether an osmotic shock is enough to induce stress granule formation. Indeed, Addition of 0.4M sucrose leads to a robust stress granule formation (**Figure 4B, C**) and we confirmed that the osmotic stress induced foci are have the same features as stress granule forming upon KRB and SCH150 as they contain also Rox8-v5 (**Figure 4D**) and polyA mRNAs (**Figure 5A**).

**Figure 5:**
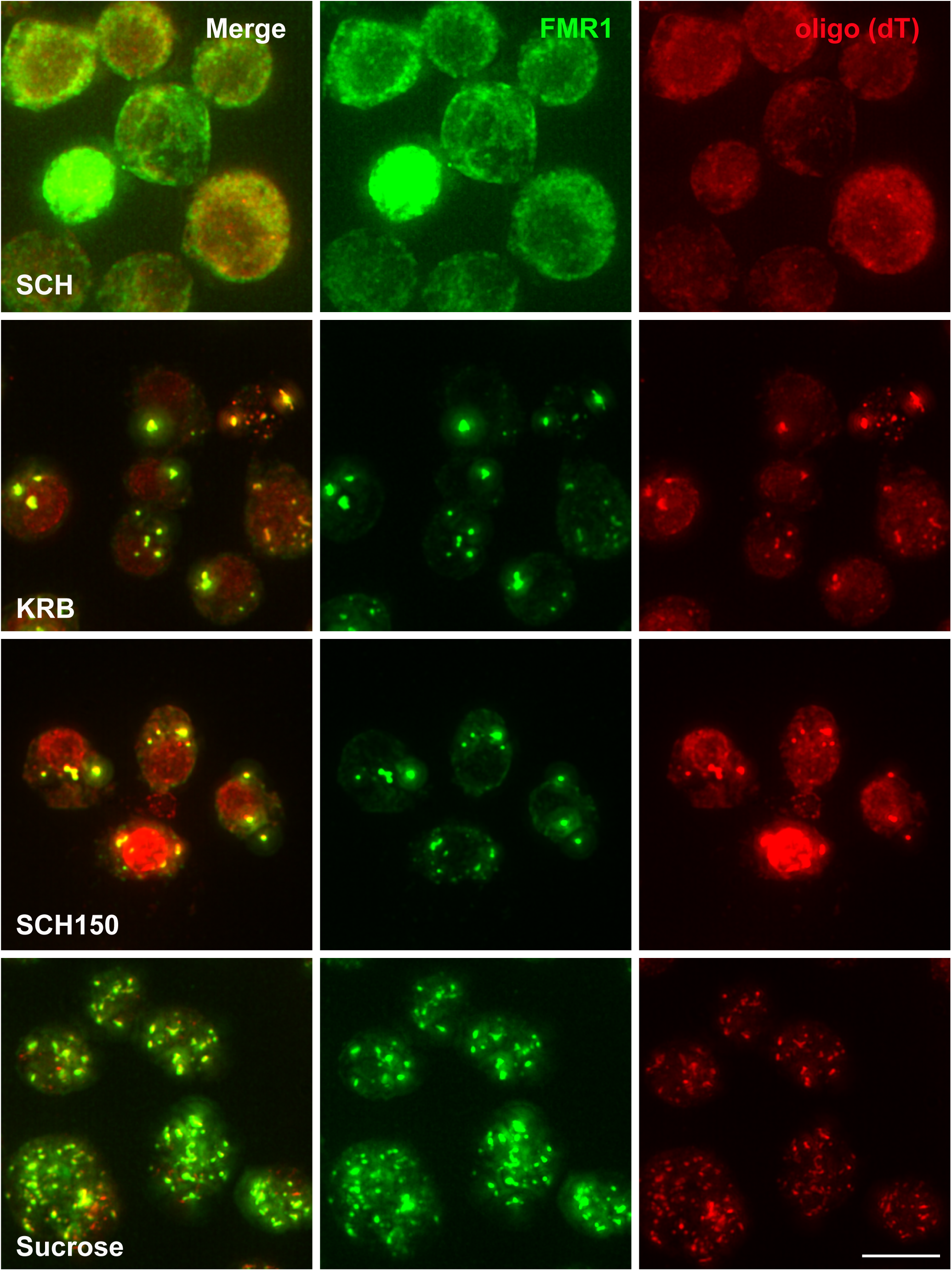
Stress granule form upon KRB and SCH150 contain polyA mRNA. Visualization of the oligo (dT) probe (red) by RNA FISH and FMRI (by IF, green marking stress granules) in cells incubated in KRB and SCH150 and SCH+sucrose. Note that stress granules are positive for polyA mRNAs. Scale bars: 10µm.

To further confirm the role of osmotic stress, we also showed that stress granule strongly formed upon addition of 150mM KCl and Na-acetate (**Figure 4B, C**). This showed that in stark contrast to Sec bodies, stress granule formation is not specific for an increase in NaCl. It also suggests that osmotic stress through an increase in extracellular salt concentration appears to be a factor leading to stress granule formation. It also suggests that KRB and SCH150 leads to this osmotic stress.

### Ca2+ increase is sufficient to drive stress granule formation

Since the addition of any salt in the medium leads to the formation of stress granule, we also tested CaCl2. This was further motivated by the results of the mass spectrometry (**Figure 6C**), showing that one of the protein whose level most increases upon KRB stress is calmodulin (**Figure 6C;** Suppl Table S1). Calmodulin binds cytoplasmic calcium and acts as signaling hub (Rajan et al., 2017).

**Figure 6:**
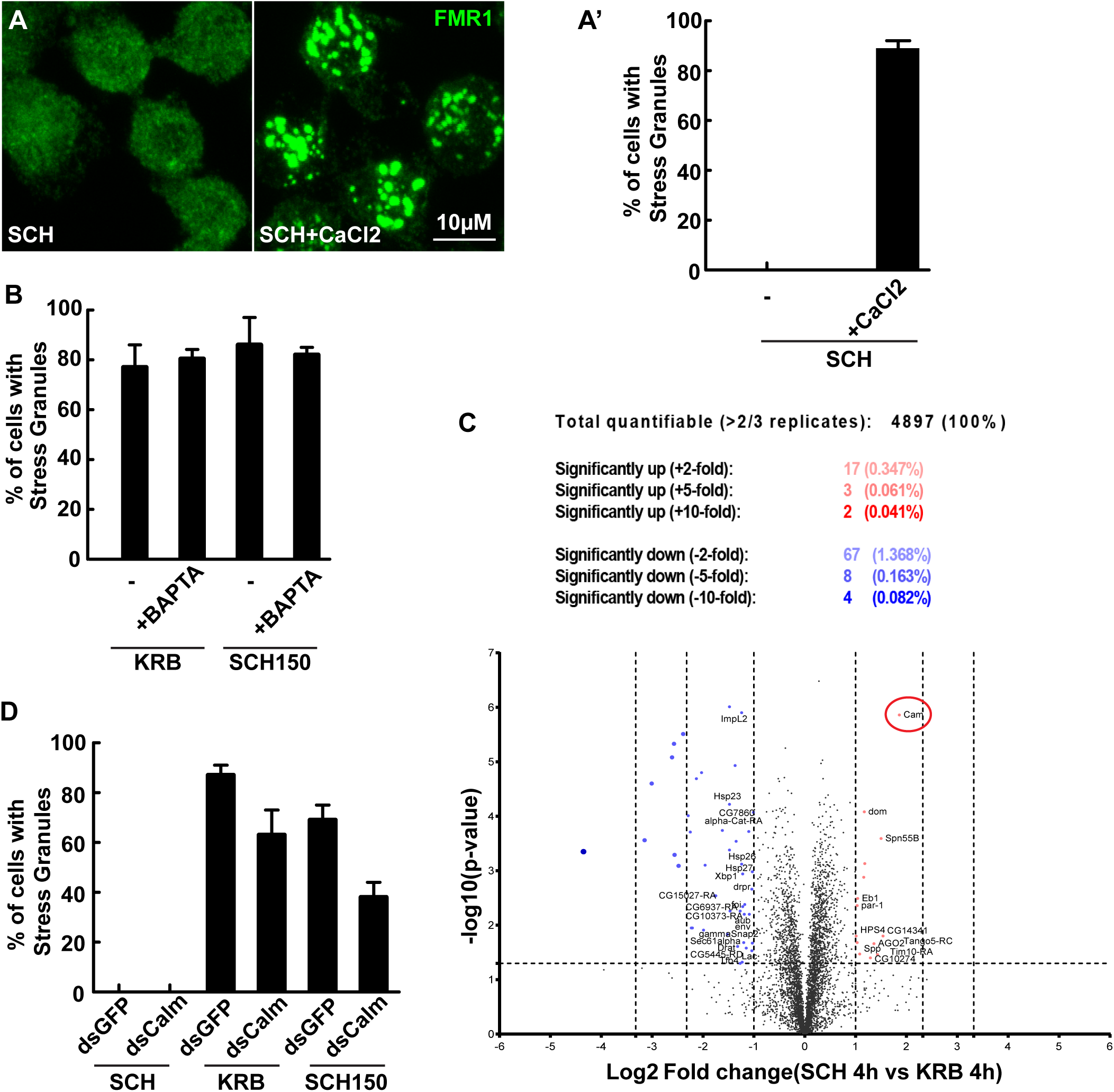
Calmodulin is involved in KRB and SCH150 driven stress granule formation. (A, A’) IF visualization of stress granule formation (marked by FMR1) in SCH and SCH with addition of CaCl_2_ (150mM) after 4 h incubation in 26°C. Quantification of stress granule formation in (A’). (B) Quantification of stress granule formation (marked by FMR1) in KRB and SCH150 with or without BAPTA-AM (BAPTA, 100µM). (C) Volcano plot of significantly regulated proteins after KRB 4 h (K) when compared to control cells kept in growing medium Schneider’s (S). 59 proteins significantly up-regulated by more than 2-fold (with paired *t*-test *p*<0.05) are indicated in red. 94 proteins significantly down-regulated by more than 2-fold (with paired *t*-test *p*<0.05) are indicated in blue. The red circle indicates Cam (calmodulin). (D) Quantification of stress granule formation in cells upon mock depletion (dsGFP) and calmodulin depletion (dsCalm) in SCH, KRB or SCH150 after 4 h incubation at 26°C. Scale bars: 10µm. Errors bars: SEM.

Addition of 150mM CaCl2 in Schneider’s strongly induced stress granule formation (**Figure 6A**), as much as NaCl (**Figure 6B**). We then asked whether a decrease in cytoplasmic calcium concentration inhibits stress granule formation. We used BAPTA-AM, a cell permeant calcium chelator, during KRB and SCH150. However, BAPTA-AM did not lead to a decrease in stress granule formation in these conditions (**Figure 6B**). Taken together, the results show that an increase of calcium in the growing medium is sufficient for stress granule formation, but that chelating it in the cytoplasm is not enough to inhibit it.

To further investigate the role of cytoplasmic calcium in stress granule formation calmodulin was depleted from S2 cells by RNAi. Depletion of calmodulin leads to 28% decrease of stress granule formation in KRB and 45% decrease in SCH150 (**Figure 6D**). Although the inhibition is only partial, these results are in line with the notion that the calmodulin, potentially via calcium is instrumental to form stress granules, and with the presence of calcium binding proteins in stress granules (Markmiller et al., 2019; Marmor-Kollet et al., 2020).

Taken together, these results show that stress granules formation in KRB and SCH150 incubated S2 cells is due to osmotic stress partially modulated by calmodulin, suggesting that calcium signaling might be involved.

## Discussion

In this study, we found stress granule formation and Sec body formation follow different pathways in Drosophila S2 cells, even though these two stress assemblies form under the same stress conditions. Our results show that stress granule formation upon those salt conditions is likely due to osmotic stress modulated by calmodulin activation.

### Osmotic stress induces stress granules along non-canonical pathways

Stress granules are phase separated non-membrane-bound reversible coalescences comprising many RNA binding proteins and RNAs (Buchan and Parker, 2009; Protter and Parker, 2016). The RNA binding protein FMR1 is one of the known stress granule markers in S2 cells. The substantial stress granule formation in salt conditions led us to ask whether those stress granules are *bonafide* stress granules, not FMR1 aggregation. First, stress granules that form in S2 cells in KRB and SCH150 contain multiple RNA binding proteins such as Rox8 (this study), Caprin and Rasputin (Aguilera-Gomez et al., 2017), although their full protein content has not been investigated as it has in mammalian cells, (Markmiller et al., 2019; Marmor-Kollet et al., 2020). Furthermore, they are fully reversible consistent with their formation through phase separation.

Osmotic stress leads to stress granule formation through non-canonical pathways that are independent of eIF2alpha phosphorylation (Anderson and Kedersha, 2002; Aulas et al., 2017), and we find here that it is also the case. Furthermore, mmammalian stress granules formed upon osmotic stress appear to contain only a subset of polyA mRNAs (Van Leeuwen and Rabouille, 2019). Here, we show that osmotic stress induces stress granule formation are positive for polyA mRNAs, albeit at a lower level than in other conditions, suggesting that they are potentially non-canonical. Whether eIF4A inhibition by RocA and PatA (Anderson and Kedersha, 2002; Aulas et al., 2017) is involved remains to be elucidated.

### Stress granule and Sec body formation in KRB and SCH150 follow different signaling pathways

Stress granules and Sec bodies form side by side and in similar time frame upon KRB and SCH150 incubation (Figure 1). However, their formation follows different pathways. Sec body formation requires the stimulation of two signaling pathways (Zhang et al., 2021). The first is ER stress via IRE1 and PERK activation. However, they do not appear to be sufficient for stress granule formation. The second pathway is high NaCl stress (addition of 150mM NaCl) via SIKs activation, which is sufficient and necessary for Sec body formation (of note, when the NaCl stress is moderate, it does not lead to Sec body formation, IRE1 and PERK activation are needed). However, SIKs activation is not necessary or sufficient for stress granule formation.

Furthermore, Sec body formation requires specific NaCl concentration and only increasing NaCl leads to a substantial formation of Sec bodies. Neither the addition of KCl, Na-acetate nor osmotic shock (sucrose) induces their formation. However, stress granule appears to be formed by the non-differential addition of salts. Addition of sucrose also leads to their formation. As we show that KRB and SCH150 do not lead to a decrease in cell diameter (no apparent shrinkage), it suggests that osmotic salt stress activates other pathways (see below). Taken together, these results suggest that the same cellular stresses induce the formation of two stress assemblies upon the activation of many pathways, a subset of them specifically leading the formation of one assembly while a different subset leads to the formation of the other. It suggests that these incubations are complex. This can also explain that, also they form in close proximity to one another near the ERES (Zacharogianni et al., 2014) and share some components Rasputin (Aguilera-Gomez et al., 2017), these two assemblies remain distinct from one another.

### ER stress can lead to stress granule formation, but it is not required upon KRB and SCH150 induced stress granule formation

It has been widely reported that stress granules are RNA-containing stress assembly which forms in response to ER stress (Goodier et al., 2007; Kimball, 2003). It is also the case for S2 cells where addition of DTT and ammonium persulfate (APS) and heat stress leads to the formation of stress granules **(Figure 7)**.

**Figure 7:**
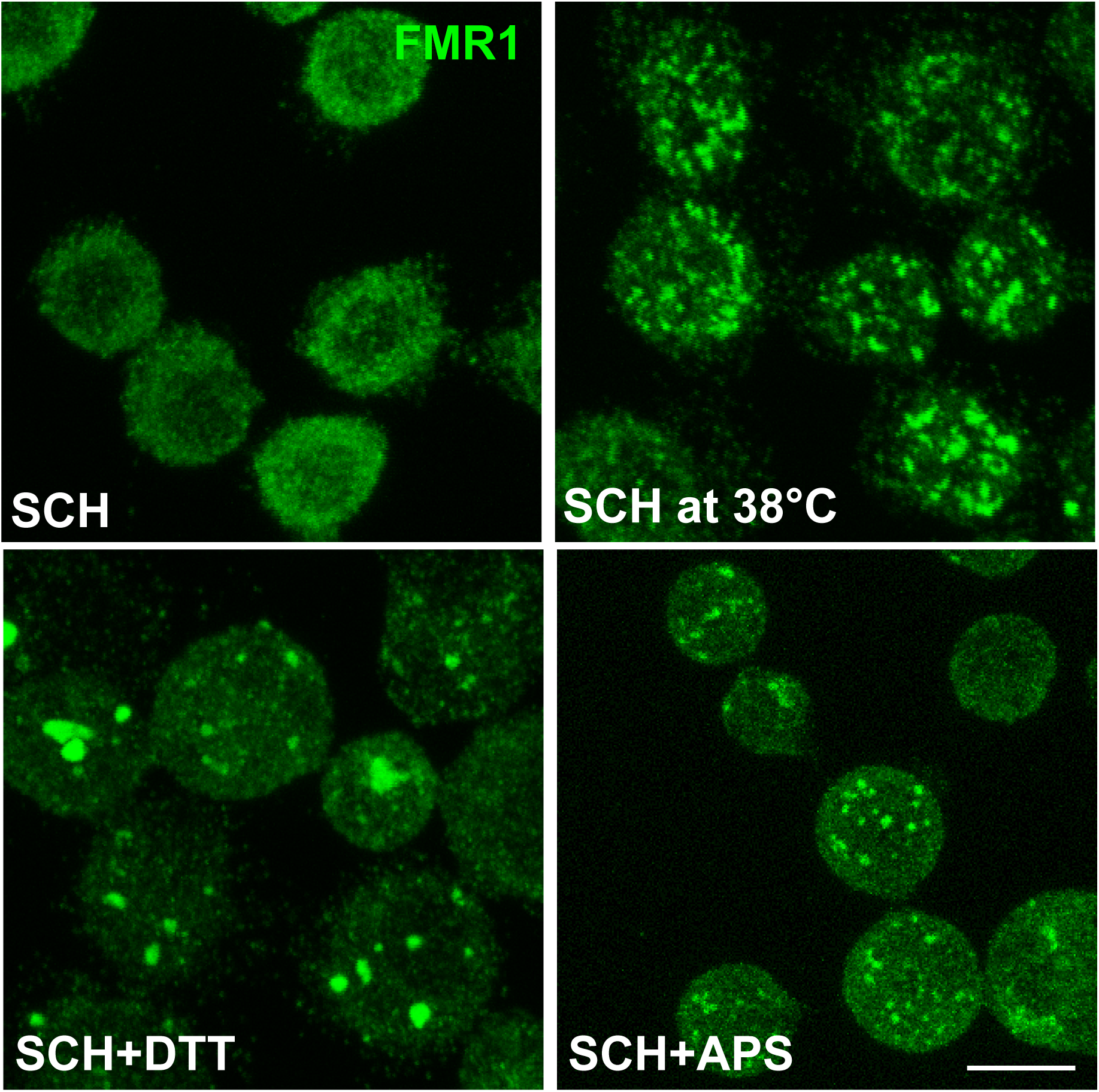
ER stress leads to stress granule formation in S2 cells. (A) IF visualization of stress granule formation (marked by FMR1) in SCH and SCH supplemented by DTT (5mM), APS (500uM) for 4 h at 26°C, and SCH for 1 h at 38°C (heat shock). Scale bars: 10µm.

This strongly suggested that ER stress of S2 cells could be instrumental to stress granule formation, at least upon KRB treatment, as we have shown that it stimulates ER stress (Zhang et al., 2021). However, neither of the ER stress inhibitors (AMG-18, 4u8C, PERKi), nor PERK and ATF6 depletion decrease stress granule formation upon KRB and SCH150. This suggests that although ER stress is sufficient to induce stress granules, ER stress kinases are not involved in KRB and SCH150 induced stress granule formation. of note, this is also the case for mTORC1 signaling. It is inhibited in KRB (Zhang et al., 2021), but its sole inhibition does not lead to stress granule formation.

### Ion level change upon osmotic stress leads to stress granule formation via calmodulin

The assessment of the intracellular Na+ concentration suggests that it increases upon KRB and SCH150 incubation. In yeast (Batiza et al., 1996; Matsumoto et al., 2002) and in plant (Tracy et al., 2008), increasing NaCl in the medium is known to lead to an increase of the cytoplasmic calcium concentration through the release of calcium from membrane-bound organelles, such as the ER, the mitochondria, the compartment of the secretory pathway and the endo-lysosomes (Medina et al., 2011). Increasing extracellular Ca^2+^ levels have been shown leads to P-body formation in yeast, in a functional calmodulin dependent manner(Kilchert et al., 2010). Here, we show that calmodulin is also involved in stress granule formation in our system, suggesting an involvement of calcium signaling. Interestingly, Ca^2+^ binding protein, including calmodulin 1 and 2 have been shown to be incorporated into mammalian stress granules (Markmiller et al., 2019; Marmor-Kollet et al., 2020). Their recruitment is in line with the finding that modulating the intracellular calcium level has an impact on stress granule formation.

Taken together, our study shows that the same cells under the same stresses form two different stress assemblies with similar kinetics by activation of multiple pathways that are differentially used. This could partially explain why the two stress assemblies remain distinct instead of completely phase separating in a single dual structure, even if they form in a close proximity (Zacharogianni et al., 2014) and share component (Aguilera-Gomez et al., 2017).

## Materials and methods

### Cell culture, KRB incubation, drug treatments and depletions by RNAi

Drosophila S2 cells (R69007, Thermo Fisher Scientific) were cultured in Schneider’s medium (Sch, S0146; Sigma) supplemented with 10% insect-tested fetal bovine serum (F4135; Sigma) at 26°C. S2 cells (between passages 5 and 18) were pelleted at 200 g in a microfuge for 3 min, washed once in fresh Schneider’s medium, and diluted to 10^6^/ml. 1 ml of cell suspension were plated per well in a 12-well plate containing coverslips. Cells were allowed to attach for 1.5 h before starting the treatment.

Amino acid starvation was performed in Krebs Ringers bicarbonate buffer (KRB) comprising 0.7 mM Na2HPO4, 1.5 mM NaH2PO4, 15 mM NaHCO3 (sodium bicarbonate), 120.7 mM NaCl, 4.53 mM KCl, 0.5 mM MgCl and 10 mM glucose at pH 7.4 as reported (Zhang et al., 2021). SCH84, SCH100 and SCH150 correspond to SCH supplemented with 84, 100 and 150 mM NaCl.

Wild-type Drosophila S2 cells were depleted by dsRNAs, as for 5 days previously described (Kondylis and Rabouille, 2003), typically leading to depletion in more than 90% of the cells. Primers for depletion used were:

dsPERK forward, 5′-TAATACGACTCACTATAGGGAGCTGGAGCTGGCTGTTTT-3′;

dsPERK reverse, 5′-TAATACGACTCACTATAGGGTACTGGCGGATATCGGCTTC-3′,

dsATF6 forward, 5′-TAATACGACTCACTATAGGGAGCGGCATGTCATAGCTGTA-3′,

dsATF6 reverse, 5′-TAATACGACTCACTATAGGGTTGACGAGAAATGCAATCCA-3′,

dsCalmodulin forward, 5’-TAATACGACTCACTATAGGGCACCTACAAAAATGGCCGA-3’,

dsCalmodulin reverse, 5’-TAATACGACTCACTATAGGGTCTTCGTAATTGACCTGACCG-3’.

### Molecular cloning and transfection

To generate pMT-Rox8-V5, Rox8 was amplified from a cDNA library made from S2 cells and cloned into pMT-V5 using the restriction enzymes KpnI and EcoRv. Primer used were: Rox8 forward: 5’-GGGATCTAGATCGGGGTACCATGGACGAGTCGCAACCG-3’, Rox8 reverse: 5’-GCCACTGTGCTGGATATCTTGGGTCTGGTATTGTGGCATCG-3’.

### Cell treatment

Drug treatment was performed on the plated cells at 26°C for 4 h incubated either in Schneider’s medium or in KRB or in SCH150 (as in (Zhang et al., 2021).

### Antibodies

For immunofluorescence, we used the rabbit polyclonal anti-Sec16 (1:800) (Ivan et al., 2008) to detect Sec16, the mouse monoclonal anti-FMR1 (1:20, deposited by Siomi, H. DSHB). Donkey anti-rabbit-IgG conjugated to Alexa Fluor 568 (1:200, A10042, Invitrogen) and a goat anti-mouse-IgG conjugated to Alexa Fluor 488 (1:200 A11001, Invitrogen) were used as secondary antibodies.

For western blotting, we used a rabbit a rabbit monoclonal antibody anti-Phospho-eIF2α (Ser51) (1:1000, 9721S, Cell Signaling) and a mouse monoclonal anti-α-tubulin (1:2500, T5168, Sigma-Aldrich) followed by anti-rabbit-IgG and mouse-IgG antibodies coupled to HRP (1:2000, NA934, NA931, GE Healthcare).

### Immunofluorescence

For immunofluorescence, cells were fixed with 4% paraformaldehyde in PBS (pH 7.4) for 20 min. Cells were then washed three times with PBS and subsequently quenched by incubation in 50 mM NH4Cl in PBS for 5 min followed by permeabilization with 0.11% Triton X-100 for 5 min. Thereafter, cells were washed three times in PBS and blocked in PBS supplemented with 0.5% fish skin gelatin (G7765, Sigma-Aldrich) for 20 min. Cells were then incubated with the primary antibody (in blocking buffer) for 25 min, washed three times with blocking buffer and incubated with the secondary antibody (in blocking buffer) coupled to a fluorescent dye for 20 min. Cells on the coverslip were washed twice in milliQ water and dried for 3 min on a tissue with cells facing up. Finally, each coverslip with cells was mounted with Prolong antifade medium (+DAPI, P36935, Invitrogen) on a microscope slide. Samples were viewed with a Leica SPE confocal microscope using a 63× oil lens and 2× zoom.

### Quantification

The quantification of stress granule formation was performed by counting the cells in which 3-5 FMR1 positive foci form (visualized by Immunofluorescence). Experiments were performed two times or more unless otherwise stated. At least four to five fields (∼25–30 cells per field) were recorded and analyzed per experiment as in (Zhang et al., 2021)

### Sodium Green Indicator assay

The stock solution of Sodium Green (5 mM in DMSO) was diluted to 5 µM in Schneider’s medium directly before usage. 10^6^ cells/ml were incubated with 5 µM Sodium Green in Schneider’s in the dark for 1h at 26°C in a 48-well plate. After incubation, the cells were washed three times with Schneider’s medium to remove excess Sodium Green. The cells were then incubated in the treatment medium and the fluorescence intensity of the dye was immediately recorded over a period of 1h at 26°C using a Spark multimode microplate reader (Tecan) with an excitation of 480 nm and an emission of 530 nm. The fluorescence intensity of five fields of cells per condition was measured every 5 min for a period of 1 h. Each experiment was performed at least three times or more. The differences in Sodium Green intensity for each condition was calculated by using the last value (1 h) minus the first value (0 h).

### RNA FISH

KRB, SCH150, 0.4M sucrose wild-type S2 cells were fixed and labeled for endogenous FMR1. After incubation with the secondary antibody, cells were washed three times with PBS and cells were post-fixed in 4% paraformaldehyde in PBS (pH 7.4) for 10 min. Following a washing three times in PBS, cells were further incubated for 5 min in 10% formamide (17899, Thermo Fisher Scientific) in DEPC-treated water. They were then incubated overnight on a droplet containing one fluorescent RNA FISH (polydT) probe [125 nM in 1% dextransulfate (D8906, Sigma-Aldrich), 10% formamide in DEPC-treated water at 37°C] in a moistened chamber to avoid drying. Cells were washed twice for 30 min with 10% formamide in DEPC-treated water and mounted with Prolong antifade medium (plus DAPI) on a microscope slide. The TMR-oligo(dT) 30× was purchased from IDT. A widefield Leica MM-AF microscope with a 100× lens was used for imaging.

### Mass Spectrometry Proteome analysis

8 million cells per condition were grown in Schneider’s and starved in KRB for 2h and 4h in 6 cm dishes at 26°C. After incubation, cells were harvested cells on ice, and cell pellets were and washed twice with ice-cold PBS. Cell material was lysed by gentle vertexing in 8M Urea in 50mM ammonium bicarbonate supplemented with 50µg/ml DNAse I (Sigma-Aldrich), 50µg/ml RNAse A (Sigma-Aldrich) and 1x complete EDTA-free protease inhibitor cocktail (Roche Diagnostics). Subsequently, the lysate was cleared by centrifugation for 1h at 18,000 x g at 15°C. Protein concentration was determined with the Bradford assay (Bio-Rad). For each sample, 20µg of total protein was reduced, alkylated and digested sequentially with Lys-C (1:100) and trypsin (1:75), and perfectionated pre-fractionated offline on C18 STAGE-tips. Peptides were eluted in 5 high-pH reversed phase fractions, with 11-80% acetonitrile. All samples were dried by vacuum centrifugation and reconstituted in 102% formic acid prior to LC-MS/MS analyses.

MS data was acquired with an UHPLC 1290 system (Agilent) coupled to a Q-Exactive HF mass spectrometer (Thermo Fischer Scientific). Peptides were trapped (Dr Maisch Reprosil C18, 3µM, 2cm x 100µM) for 5min in solvent A (0.1% formic acid in water) before being separated on an analytical column (Agilent Poroshell, EC-C18, 2.7µM, 50cm x 75µM). Solvent B consisted of 0.1% formic acid in 80% acetonitrile. The mass spectrometer operated in data-dependent mode. Full scan MS spectra from m/z 375 – 1600 were acquired at a resolution of 60,000 to a target value of 3×106 or a maximum injection time of 20ms. The top 15 most intense precursors with a charge state of 2+ to 5+ were chosen for fragmentation. HCD fragmentation was performed at 27% normalized collision energy on selected precursors with 16s dynamic exclusion at a 1.4m/z isolation window after accumulation to 1×10^5^ ions or a maximum injection time of 50ms. Tandem mass spectrometry (MS/MS) spectra were acquired at a resolution of 15,000.

## Acknowledgments

We thank Sem Brussee for the calmodulin depletion experiment, and the Hubrecht Imaging Center for support with microscopy. We acknowledge Genentech for providing us with the IRE1 inhibitor AMG18.

